# Systemic antibody-oligonucleotide delivery to the central nervous system ameliorates mouse models of spinal muscular atrophy

**DOI:** 10.1101/2021.07.29.454272

**Authors:** Suzan M Hammond, Frank Abendroth, Larissa Goli, Matthew Burrell, George Thom, Ian Gurrell, Jessica Stoodley, Nina Ahlskog, Michael J Gait, Matthew J A Wood, Carl Webster

## Abstract

Antisense oligonucleotides (ASOs) have emerged as one of the most innovative new genetic drug modalities, however, the high molecular weight limits their bioavailability for otherwise treatable neurological disorders. We investigated conjugation of ASOs to an antibody against the murine transferrin receptor (TfR), 8D3_130_, and evaluated it via systemic administration in mouse models of the neurodegenerative disease, spinal muscular atrophy (SMA). SMA like several other neurological and neuromuscular diseases, is treatable with single-stranded ASOs, inducing splice modulation of the survival motor neuron 2 (*SMN2*) gene. Administration of 8D3_130_-ASO conjugate resulted in bioavailability of 2.7% of the injected dose in brain. Additionally, 8D3_130_-ASO yielded therapeutically high levels of *SMN2* splicing in the central nervous system of mildly affected adult SMA mice and resulted in extended survival of severe SMA mice. Systemic delivery of nucleic acid therapies with brain targeting antibodies offers powerful translational potential for future treatments of neuromuscular and neurodegenerative diseases.

## Introduction

RNA targeted therapeutics are an emerging form of therapy amendable to treating multiple neurodegenerative and neuromuscular diseases, many of which are currently without a therapeutic intervention. Antisense oligonucleotides (ASOs) are synthetic single-stranded oligonucleotides or oligonucleotide analogues designed to bind to RNAs, either messenger or noncoding RNA, through Watson-Crick base pairing to modulate their function. ASOs act either through steric blockage of cis regulatory elements on mRNAs or by inducing RNase H1 mediated degradation of the targeted RNA(*1*). There are currently 8 FDA approved single-stranded ASO drugs and many more under pre-clinical investigation(*2*). One of these, nusinersen, has been approved for the treatment of spinal muscular atrophy (SMA). SMA is a neurodegenerative disease characterized by the loss of lower motor neurons of the spinal cord and subsequent skeletal muscle atrophy. SMA is caused by a reduced level of survival motor neuron (SMN) protein due to mutations in the *SMN1* gene. Humans carry a redundant paralogue of *SMN1*, called *SMN2*, which is unaffected in the majority of SMA patients. However, *SMN2* has two main splice variants; full-length *SMN2* (*FLSMN2)* mRNA (~10%) yields functional SMN protein, while *Δ7SMN* (~90%) generates an unstable truncated SMN protein that is typically degraded(*3–6*). ASOs bind to the intron splice suppressor N1 (ISS-N1) of *SMN2* exon 7 pre-mRNA to block splicing factor binding thus increasing the probability of exon 7 incorporation and thereby increasing the level of mature *FLSMN2* mRNA. The clinically successful ASO therapy for SMA (Spinraza/nusinersen) has received worldwide regulatory approval (*7*).

Systemic ASO therapy of neurodegenerative diseases is made challenging by the need to pass through the blood-brain barrier (BBB) and blood spinal cord barrier (BSCB). On their own, systemically administered ASOs are readily accessible in tissues such as liver but only very moderately bioavailable in peripheral tissues such as skeletal muscle. Advanced modifications to the backbone and sugar moieties of ASOs have improved tissue uptake but have yet to achieve significant BBB penetration for exposure in brain and spinal cord. As such, ASO compounds approved (including nusinersen) or in development for neurodegenerative disease typically circumvent the BBB/BSCB via local administration directly into the circulating cerebrospinal fluid compartment via the intrathecal route(*8*). However, the intrathecal route is sub-optimal, typically resulting in lower levels of ASO exposure in higher spinal cord and brain regions, and this route is precluded in certain SMA patients such as those with spinal column abnormalities(*9*). Additionally, following intrathecal administration, low levels of ASO have been reported outside the CNS, for example in important target tissues such as skeletal muscle and liver, both known to be affected in SMA. Therefore, a systemically administered ASO delivery system with biological activity in brain and spinal cord as well as in peripheral tissues, is required to facilitate optimal treatment in all SMA patients.

The transferrin receptor (TfR) is the most widely studied pathway for transport of antibody-based drugs across the BBB, blood spinal cord barrier (BSCB) and choroid plexus(*10–13*). TfR is expressed on the luminal side of brain capillary endothelial cells where it binds transferrin and traffics iron into the parenchyma. Anti-TfR antibodies and antibody fragments have successfully used this pathway to deliver drug cargoes into the brain parenchyma(*10, 12, 14, 15*). However, to achieve sufficient therapeutic levels of drug exposure within the brain, precision engineering of the binding affinity to TfR is required. Antibodies with a high affinity for TfR preferentially accumulate in brain capillaries, but do not release efficiently to the abluminal side(*16–20*). Elevating the levels of antibody accumulation in the brain parenchyma requires engineering to optimise affinity, introduce pH-dependent binding, and/or create monovalent binding(*20–22*). The anti-mouse TfR monoclonal antibody, 8D3130, was designed to bind specifically to mouse (mTfR) at sufficiently low affinity to permit BBB transcytosis by releasing it from the mTfR once exposed to the brain parenchyma(*15, 23*).

Our previous work has demonstrated the ability of cell-penetrating peptides to deliver morpholino phosphorodiamidate oligonucleotides (PMOs) across the BBB and BSCB at therapeutically relevant doses(*24*). Here we report that systemic delivery of PMOs directly conjugated to the anti-mouse TfR monoclonal antibody 8D3_130_ yields even greater bioavailability in CNS (2.7% of injectable dose in the brain and 2.3% in the spinal cord). This induces high expression of full-length *SMN2* mRNA and SMN protein in the CNS and in peripheral tissues of an adult mild SMA mouse, and rescuing survival of severely affected SMA mice. This work provides a way forward for systemic ASO treatment in SMA and other neurodegenerative diseases treatable with ASOs including HD, ALS, SCA2 and tauopathies such as Alzheimer’s disease(*25*).

## Results

### Synthesis and purification of anti-TfR antibody-PMO conjugates

The intronic splicing silencer N1 (ISS*-*N1) is a regulatory element in intron 7 of the human *SMN2* (*hSMN2*) gene. It is the most promising target for modulation of *SMN2* splicing since masking it with a fully modified ASO favours inclusion of exon 7(*26–28*). We synthesized a 25-mer PMO targeting ISS-N1 and directly conjugated it to a short maleimide functionalized peptide linker (Mal-C3-FB[RB]_6_, B = beta-alanine; Fig. 1A). The maleimide-functionalized PMO was then conjugated to the heavy chain of either a low affinity anti-mouse TfR (8D3_130_) or a negative isotype-control antibody against nitrophenol (NIP228)(*15*) (Fig. 1A). The PMO-antibody conjugates were purified via size exclusion chromatography to remove any unreacted maleimide-functionalized PMO and antibody. Analysis of purified antibody-conjugates by MALDI-TOF mass spectrometry revealed a fully modified product with a drug to antibody ratio of 2, i.e. the incorporation of 2 PMOs per antibody, which correspond to an overall mass shift of 20.4 kDa (Fig. 1B, S1 and S2).

**Figure 1:**
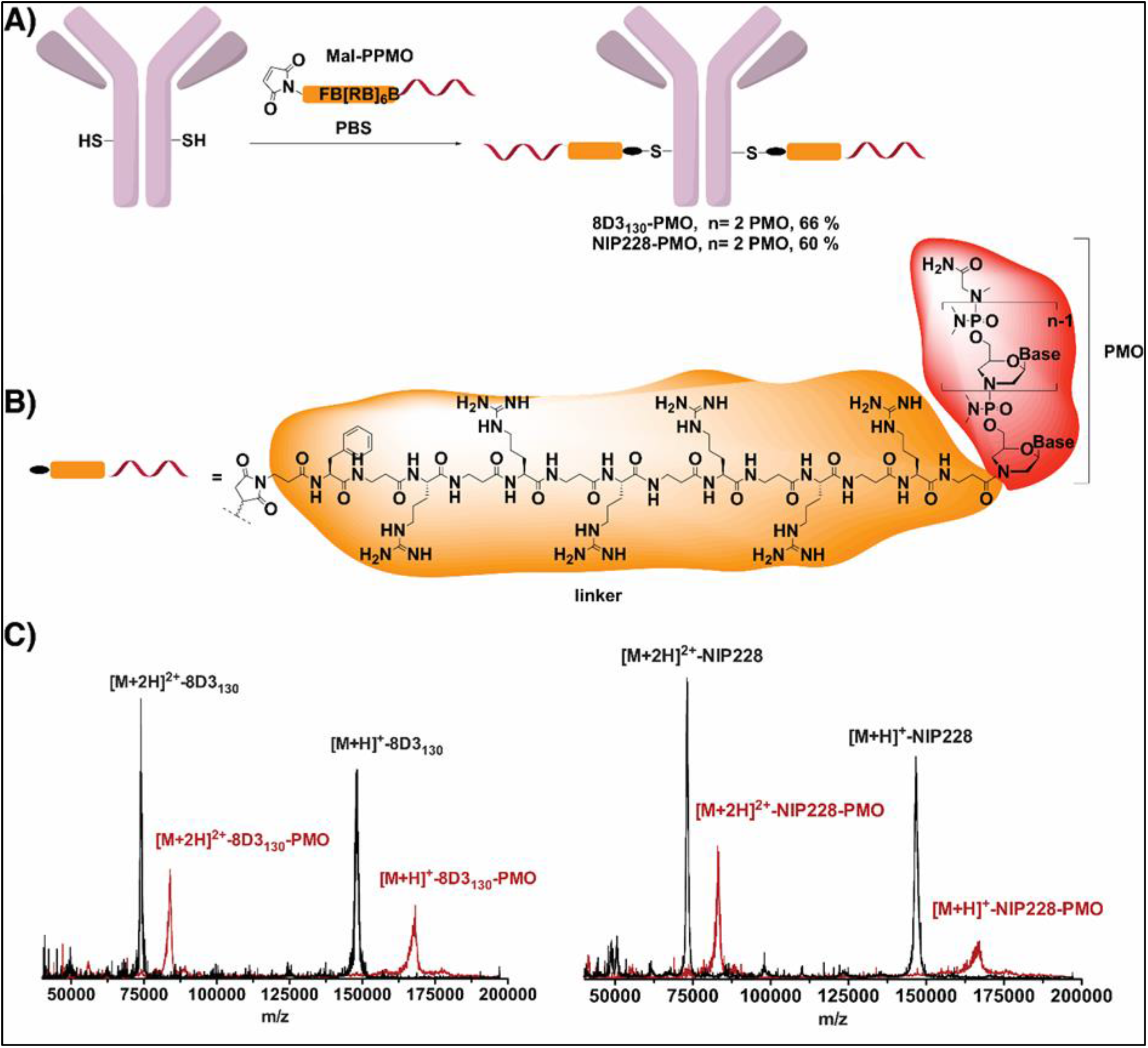
Synthesis of Antibody-PMO Conjugates. (A) Schematic of conjugation synthesis between 25 mer phosphorodiamidate morpholino oligo (PMO) targeting the ISS-N1 site of the SMN2 gene to the free thiol group of a solvent exposed engineered cysteine residue in the CH2 domain of the heavy chain of either a low affinity mouse transferrin receptor (8D3_130_) or a negative control antibody (NIP228). (B) Chemistry of linker between PMO and antibodies. (C) Maldi-Tof spectra of the unmodified antibodies (black trace) and the purified Antibody-PMO-conjugates (red traces) for the 8D3_130_ (left) and NIP228 antibodies (right).

### PMO conjugation alters the pharmacokinetic properties of the 8D3_130_ antibody

To determine the pharmacokinetic (PK) properties of the antibody-PMO conjugates 8D3_130_– PMO and NIP228-PMO, and whether these altered the plasma, brain and spinal cord exposure over antibody alone, we conducted a plasma PK, brain and spinal cord exposure study *in vivo*. Adult C57BL/6J mice were administered with a single intravenous (IV) dose of 20 mg/kg of 8D3_130_ (with or without PMO) or isotype control antibody NIP228 (with or without PMO) and blood, brain and spinal cord homogenate samples were studied at regular time intervals over a one-week period. A MesoScale Discovery (MSD) Universal hIgG capture and detection assay for 8D3_130_ and NIP228 antibodies was used to determine exposure parameters.

Brain and spinal cord samples were collected at 4, 24, 96 and 168 hours post-administration and homogenates processed for analysis *via* MSD assay (Fig. 2, A and B). The MSD plate-based sandwich immunoassay was formatted whereby the anti-human IgG capture antibody bound to sample hIgG (± PMO), and a specific detection antibody labelled with SULFO-TAG emitted light on electrochemical stimulation. The levels of hIgG (± PMO) in plasma, brain and spinal cord were quantified by reference to standard curves generated using calibrator samples with a four-parameter nonlinear regression model. At 24 h post-administration, the C_max_ of 8D3_130_ (unconjugated and conjugated) in the brain (Fig. 2A), was significantly higher than NIP228 (10-fold and 6-fold for conjugated and unconjugated, respectively; Table 1). At one week, the total exposure to the drug, measured by the area under the curve (AUC_last_) of 8D3_130_ and 8D3_130_-PMO was 8- and 5-fold higher, respectively, than NIP228-PMO (Table 1). Brain and spinal cord measurements for NIP228-PMO were below the lower limit of quantification (LLoQ) for the assay for all time points and are not presented here.

**Figure 2:**
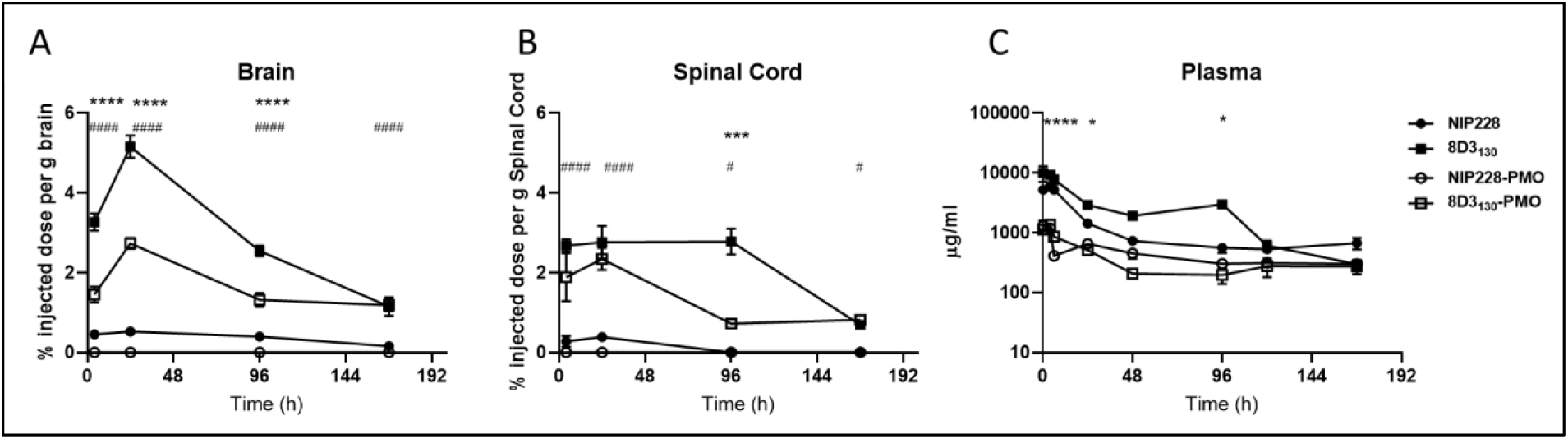
Pharmacokinetics of antibody and antibody-PMO conjugates in mouse. Plasma, brain and spinal cord exposure following a 20 mg/kg dose of the anti-TfR antibody, 8D3_130_ and isotype control antibody, NIP228; unconjugated and conjugated to PMO oligonucleotide in a Universal PK assay. (A) Brain exposure as a measure of % injected dose per gram of brain. Statistical significance (****) shown for 8D3_130_ 20 mg/kg vs 8D3_130_-PMO 20 mg/kg for first three time points. Statistical significance (^####^) shown for NIP228-PMO 20 mg/kg vs 8D3_130_-PMO 20 mg/kg at all time points. (B) Spinal cord exposure as a measure of % injected dose per gram of spinal cord. Statistical significance (***) shown for 8D3_130_ 20 mg/kg vs 8D3_130_-PMO 20 mg/kg at 96 h time point. Statistical significance (^####^) shown for NIP228-PMO 20 mg/kg vs 8D3_130_-PMO 20 mg/kg at first two time points with lower statistical significance (^#^) at the last two time points. (c) Plasma PK of antibodies with or without PMO over a one-week period. Statistical significance (****) shown for 8D3_130_ 20 mg/kg vs 8D3_130_-PMO 20 mg/kg for the first three time points with lower significance (*) for 24 and 96 h time-points. No statistical significance shown for NIP228-PMO 20 mg/kg vs 8D3_130_-PMO 20 mg/kg at any time point. Statistical significance (representative P values) for exposure in brain, spinal cord and plasma between 8D3_130_ 20 mg/kg vs 8D3_130_-PMO 20 mg/kg (*) and NIP228-PMO 20 mg/kg vs 8D3_130_-PMO 20 mg/kg (#) at all time points tested. Statistical analysis was performed in GraphPad Prism. Data shown as the mean +/− standard error of the mean, n = 3-4 per group. Statistical significance shown using 2-way analysis of variance, where appropriate, were made using Tukey test. *P, <0.05; **P, <0.01; ***P, <0.001; ****P, <0.0001; ^#^P, <0.05; ^##^P, <0.01; ^###^P, <0.001; ^####^P, <0.0001; ns, not significant.

**Table 1.** Summary of brain and spinal cord exposure of 8D3_130_, 8D3_130_-PMO, and NIP228 in C57BL/6J mice following intravenous administration.

| Construct | Brain |  |  | Spinal Cord |  |  |
| --- | --- | --- | --- | --- | --- | --- |
| | T<sub>max</sub><br>(h) | C<sub>max</sub><br>(μg/g) | AUC<sub>last</sub><br>(h*μg/g) | T<sub>max</sub><br>(h) | C<sub>max</sub><br>(μg/g) | AUC<sub>last</sub><br>(h*μg/g) |
| 8D3<sub>130</sub> | 24 | 22.6 | 2180 | 24 | 12.6 | 1460 |
| 8D3<sub>130</sub>-PMO | 24 | 13 | 1330 | 24 | 10.5 | 614 |
| NIP228 | 24 | 2.29 | 263 | 24 | 1.79 | 28.4 |

In the spinal cord, C_max_ at 24 h of 8D3_130_ (unconjugated and conjugated) was significantly higher than either NIP228 or NIP228-PMO (7-fold and 6-fold for conjugated and unconjugated, respectively; Table 1). AUC_last_ of 8D3_130_ and 8D3_130_-PMO was an impressive 51- and 22-fold higher, respectively, than for NIP228 (Table 1). Lower AUC for both 8D3_130_-PMO and NIP228-PMO are likely explained by the increased clearance of the conjugated forms (discussed below).

At its highest absorption (C_max_), 8D3_130_ was 8-fold higher and NIP228 was 4-fold higher than their respective PMO-conjugated antibodies. The addition of PMO correlated negatively with clearance (Cl (ml/h/kg)) data revealing a 4- and 2.5-fold higher clearance with PMO compared to 8D3_130_ and NIP228 without PMO (Table 2). Unconjugated 8D3_130_ had significantly higher total plasma exposure (AUC_last_, the area under the plasma concentration-time curve from time zero to the last measured concentration) than 8D3_130_-PMO during the first 4 days post-injection (Fig. 2C and Table 2). One week post-injection, the total plasma exposure (AUC_last_) was 7- and 3-fold higher for 8D3_130_ and NIP228 compared with their respective PMO-conjugated antibodies (Table 2).

**Table 2.** Summary of plasma exposure of 8D3_130_ and NIP228 with and without PMO in C57BL/6J mice following intravenous administration.

| Construct | C<sub>max</sub><br>(ug/ml) | AUC<sub>last</sub><br>(h*ug/ml) | CI<br>(ml/h/kg) |
| --- | --- | --- | --- |
| 8D3<sub>130</sub> | 9380 | 350000 | 0.054 |
| 8D3<sub>130</sub>-PMO | 1230 | 51800 | 0.240 |
| NIP228 | 5230 | 175000 | 0.078 |
| NIP228-PMO | 1330 | 69600 | 0.193 |

### Dose-dependent SMN2 upregulation in brain and spinal cord by 8D_3130_-PMO

The SMA-like *hSMN2* mouse strain (FVB.Cg-*Smn1^tm1Hung^* Tg(SMN2)2Hung/J) carries the entire human SMN2 gene, lives to adulthood, and has an unaltered BBB and BSCB(*29, 30*). This strain is therefore an ideal adult mouse model in which to test the biodistribution and biochemical efficacy of nucleic acid drug compounds that regulate *SMN2* gene expression and splicing. *SMN2* transgenic mice were given a single IV administration of escalating doses from 10-100 mg/kg of 8D3_130_-PMO or control NIP228-PMO. A matched volume of 5 ul/g saline administration was used for untreated controls. Each dosage group had n = 5 mice per group with exception of 20 mg/kg dose where n = 4 mice. Tissues were harvested 7 days post administration and analysed for *SMN* mRNA splicing by qRT-PCR (10, 20, 50, 80, and 100 mg/kg) and SMN protein (by Western blot; 10, 50, 100 mg/kg) (Fig. 3 and Fig. 4).

**Figure 3:**
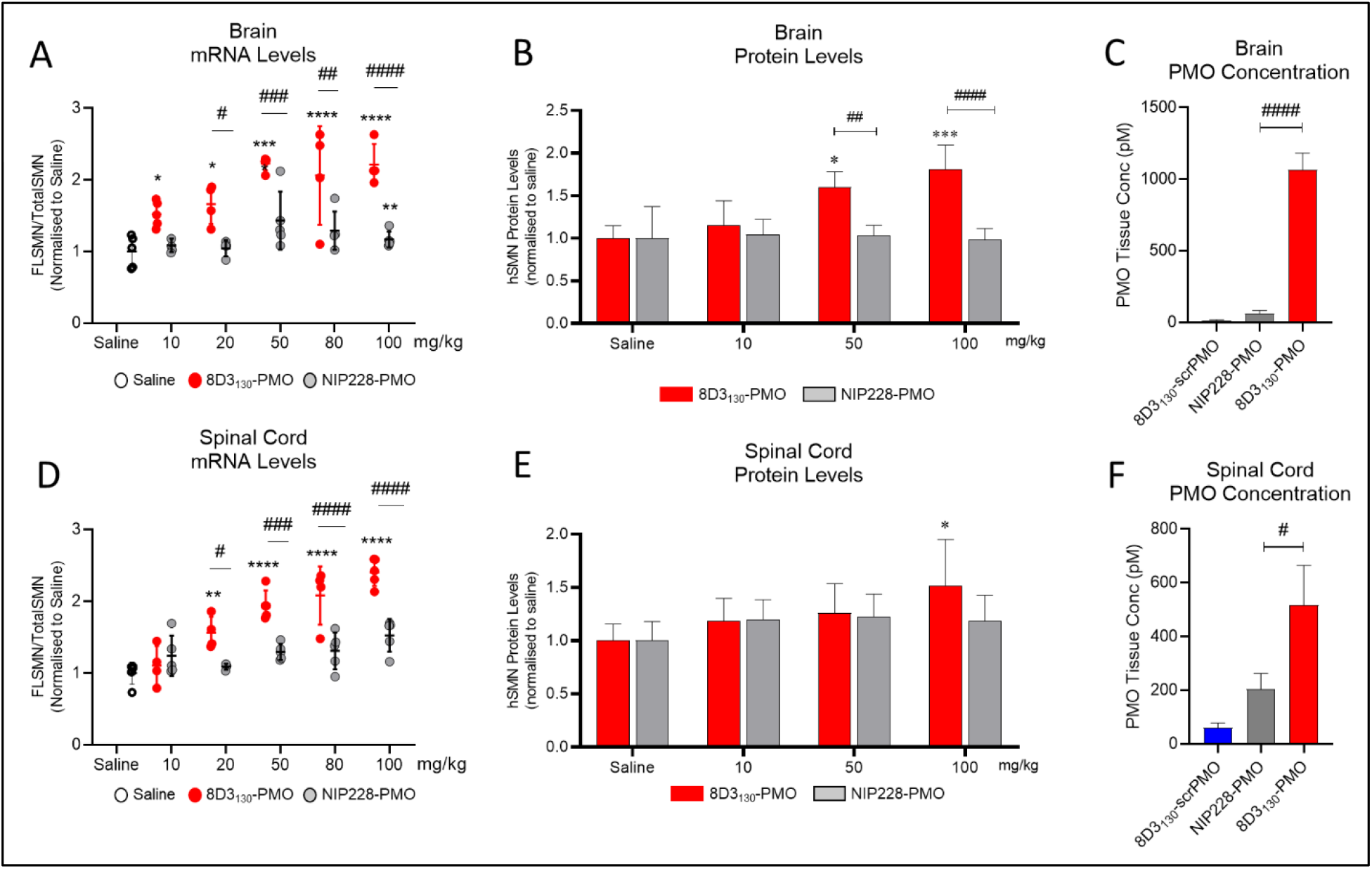
*In vivo* activity and concentration of antibody-PMOs in central nervous system of adult transgenic mice bearing the human *SMN2* gene. Tail vein administration of 10-100 mg/kg 8D3_130_-PMO and NIP228-PMO were given at 8 weeks of age and tissues harvested 7 days post-administration. Splice switching activity of the compounds compared to saline treatment on the human SMN2 transgene was analysed via qPCR and western blots (mean ± S.E.M.). qRT-PCR (A) and Western analysis (B) of brain shows distinct splicing activity between transferrin receptor targeted 8D3_130_-PMO and NIP228-PMO treatment groups, indicating transferrin receptor specific activity. (C) PMO concentration as determined by ELISA. 8D3_130_-ScrPMO is an 8D3130-PMO conjugate (with scrambled PMO sequence) used as negative control. qRT-PCR (D) and Western analysis (E) of spinal cord shows high levels of activity with less specificity as brain. (F) PMO concentration of 8D3_130_-PMO much greater than controls NIP228-PMO or 8D3_130_-scrPMO. Statistical significance (representative P values) between 8D3_130_-PMO vs Saline (*) and 8D3_130_-PMO vs NIP228-PMO (#) was performed in GraphPad Prism. Data shown as the mean ± standard deviation, n = 5-6 per group. qRT-PCR and Westerns were analysed with 2-way ANOVA corrected for multiple comparisons using Dunnett Test. ELISA for PMO concentration was analysed with 1 Way ANOVA corrected for multiple comparisons using Tukey Test. P values adjusted to account for each comparison, confidence level 0.95%. *p, <0.05; **p, <0.005; ***p, <0.0005; ****p, <0.0001; ^#^p, <0.05; ^##^p, <0.005; ^###^p, <0.0005; ^####^p, <0.0001; ns, not significant.

**Figure 4:**
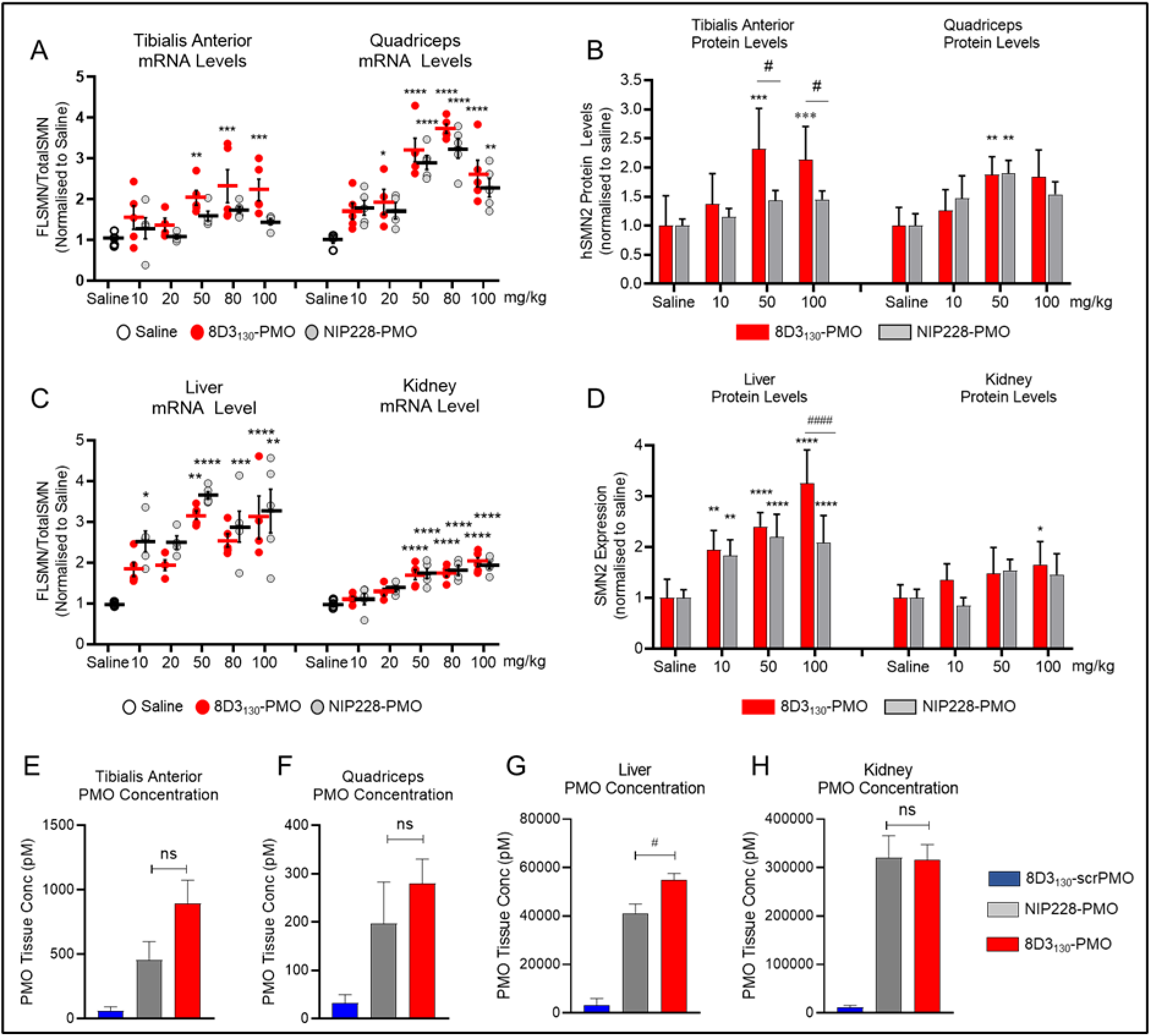
*In vivo* activity and concentration of antibody-PMOs in peripheral tissues of adult SMN2 Transgenic mice. Tail vein administration of 10-100 mg/kg 8D3_130_-PMO and NIP228-PMO were given at 8 weeks of age and tissues harvested 7 days post-administration. Splice switching activity of the compounds compared to saline treatment on the human SMN2 transgene was analysed via qPCR and western blots (mean ± S.D.). qRT-PCR (A) and Western analysis (B) of skeletal muscles tibialis anterior and quadriceps shows nearly equal activity of both 8D3_130_-PMO and NIP228-PMO with distinct splicing activity between transferrin receptor targeted 8D3_130_-PMO and NIP228-PMO treatment groups only in tibialis anterior SMN protein levels. qRT-PCR (C) and Western analysis (D) of liver and kidney tissues. Liver shows correlative activity with increasing dose administration. However, distinction between 8D3_130_-PMO and NIP228-PMO is only observable with 100 mg/kg dosing in SMN2 protein expression. PMO concentration as determined by ELISA in (E) tibialis anterior, (F) quadriceps, (G) liver and (H) kidney. Distinct uptake between 8D3_130_-PMO and NIP228-PMO was only observable in liver. All other tissues indicated similar uptake in both compounds. 8D3_130_-ScrPMO, with scrambled PMO sequence, is used as negative control. Statistical significance (representative P values) between 8D3_130_-PMO vs Saline (*) and 8D3_130_-PMO vs NIP228-PMO (#) was performed in GraphPad Prism. Data shown as the mean +/− standard deviation, n = 5-6 per group. qRT-PCR and Westerns were analysed with 2 Way ANOVA corrected for multiple comparisons using Dunnett Test. ELISA for PMO concentration was analysed with 1 Way ANOVA corrected for multiple comparisons using Tukey Test. P values adjusted to account for each comparison, confidence level 0.95%. *p<0.05, **p<0.005, ***p<0.0005, ****p<0.0001, ^#^p, <0.05; ^##^p, <0.005; ^###^p, <0.0005; ^####^p, <0.0001; ns, not significant.

*FLSMN2* mRNA (exon 7) was normalised to total SMN (qRT-PCR expression of exons 1-2a), and plotted as a fold change over saline treated expression (Fig. 3, A and D). At the lowest dose of 10 mg/kg, *FLSMN2* expression in the brain was statistically greater than saline treated mice (1.52 ± 0.08 vs 1.0 ± 0.098). A maximum average expression of 2.213 ± 0.145 fold expression of the norm (*FLSMN2* under saline treatment) was observed at 50 mg/kg dose. This activity was reflected in Western blots of SMN protein (Fig. 3B). A 50 mg/kg and 100 mg/kg administration of 8D3_130_-PMO yielded 1.6 ± 0.089 and 1.8 ± 0.128 fold change in SMN protein expression over saline treatment, respectively. To date, this is the highest level of expression published in the CNS of an SMA adult mouse model following systemic administrations of any drug. The high activity was also notable in the spinal cord. *FLSMN2* mRNA analysis of the thoracic region of the spinal cord yielded the highest fold change at 80 mg/kg (2.0 ± 0.2) and 100 mg/kg (2.4 ± 0.086) compared to saline treated controls (Fig. 3D). Protein was extracted from the cervical region of the spinal cord. At the 100 mg/kg dosing, SMN2 protein expression was significantly increased over saline treatment (1.52 ± 0.2 vs 1.0 ± 0.07), p = 0.009 (Fig. 3E). In contrast, NIP228-PMO treated tissues were not found to be significantly changed from saline treatment.

To determine the tissue concentration of PMO, a single IV dose of 50 mg/kg (n=6 per group) was chosen given its high level of activity. Seven days post-administration, tissues were collected and the concentration of PMO was determined via ELISA (*31*) (Fig. 3, C and F). A scrambled PMO conjugated to 8D3_130_ was used to control for aberrant probe binding. The tissue concentration of PMO delivered by 8D3_130_ within the brain (1066 ± 282 pM) and spinal cord (517 ± 36 pM) was significantly greater than PMO delivered by NIP228 (63 ± 53 pM and 203.5 ± 146 respectively).

Enhanced uptake and activity were also notable in the peripheral tissues. Neuromuscular diseases typically require therapies to target skeletal muscles. SMA in particular has also been shown to affect the liver and kidney tissues (*32–34*). Therefore, a potential advantage of a systemically administered drug treatment would be its activity within these tissues. We analysed the activity of 8D3_130_-PMO and NIP228-PMO in adult SMA mice. As above, mice were treated with single IV administration ranging from 10-100mg/kg antibody-PMO and tissues collected 7 days post-administration. Two skeletal muscles of the hindlimb, tibialis anterior (TA) and quadriceps (Quad) were selected for evaluation of *FLSMN2* mRNA and protein expression changes over saline treated tissues (Fig. 4, A and B). At the 20 mg/kg dose, 8D3_130_-PMO yielded a 2-fold enhanced expression over saline treated tissues (1.9 ± 0.6 vs 1.0 ± 0.17 fold change, respectively). The highest level of mRNA expression was achieved at the 50mg/kg and 80 mg/kg doses (3.2 ± 0.7 and 3.7 ± 0.3 fold change, respectively), indicating a plateau in efficacy at higher doses. High levels of expression and eventual plateau was also notable in the TA for both mRNA and protein (Fig. 4, A and B). The control NIP228-PMO also yielded high levels of activity in Quad but not in the TA for both mRNA (Quad: 3.2 ± 0.5 fold change and TA: 1.7 ± 0.17 fold change) and protein (Quad: 1.9 ± 0.2 fold change and TA: 1.43 ± 0.17 fold change)(Fig. 4, A and B). These results indicate a difference in activity that could possibly be explained by differences in muscle fibre composition between different skeletal muscles.

Both 8D3_130_-PMO and NIP228-PMO were active in the liver and kidney (Fig. 4, C and D). *FLSMN2* mRNA expression in liver was highest at 50 mg/kg for both 8D3_130_-PMO (3.17 ± 3.6 fold change) and NIP228-PMO (3.6 ± 0.2 fold change). There was no significant difference between the two antibody-PMOs in mRNA expression, however, the protein expression was greater in 8D3_130_-PMO treatment at 100mg/kg (3.2 ± 0.6 fold change) than NIP228-PMO (2.1 ± 0.5 fold change), p < 0.0001 (Fig. 4B). In the kidney there appeared to be a plateau in activity from 50 mg/kg both at the mRNA and protein levels (Fig. 4, C and D). Kidney expression of *FLSMN* mRNA (Fig. 3C) and protein (Fig. 3F) was low for all doses. Low levels of renal uptake are expected because the molecular mass of 8D3_130_-PMO (~170 kDa) is far greater than the 60 kDa filtration cut-off for kidney filtration. PMO concentration in the tissues reflected the molecular effects: both 8D3_130_ and NIP228 delivered high levels of PMO into each tissue (Fig. 4, E to H). However, only in the liver was there a significant difference between the concentration of 8D3_130_-PMO vs NIP228-PMO (p = 0.02). It is understood that high levels of circulating mAb saturate the IgG binding FcRn recycling pathway and increase the clearance of mAbs from tissues(*35, 36*).

### 8D3_130_-PMO localisation throughout the brain

Previous studies have shown that anti-TfR antibodies typically require 24 hours to translocate through the endothelia of the BBB or BSCB to enter the parenchyma of the brain and spinal cord(*20*). To observe distribution in the brain we dosed *hSMN2* transgenic mice with 50 mg/kg 8D3_130_-PMO or NIP228-PMO (n=3 per group) and perfused animals 24 hours later. Cryosections of the brain were immunohistochemically stained for 8D3_130_-PMO and NIP228-PMO using an Alexa Fluor 488 goat anti-human IgG secondary antibody (Fig. 5, A and B). Whole brain images revealed a fluorescent signal for 8D3_130_-PMO throughout the brain. The strongest staining of 8D3_130_-PMO was observed in the thalamus, pons and cerebellum regions of the brain. In contrast, NIP228-PMO was not observable in the brain.

**Figure 5:**
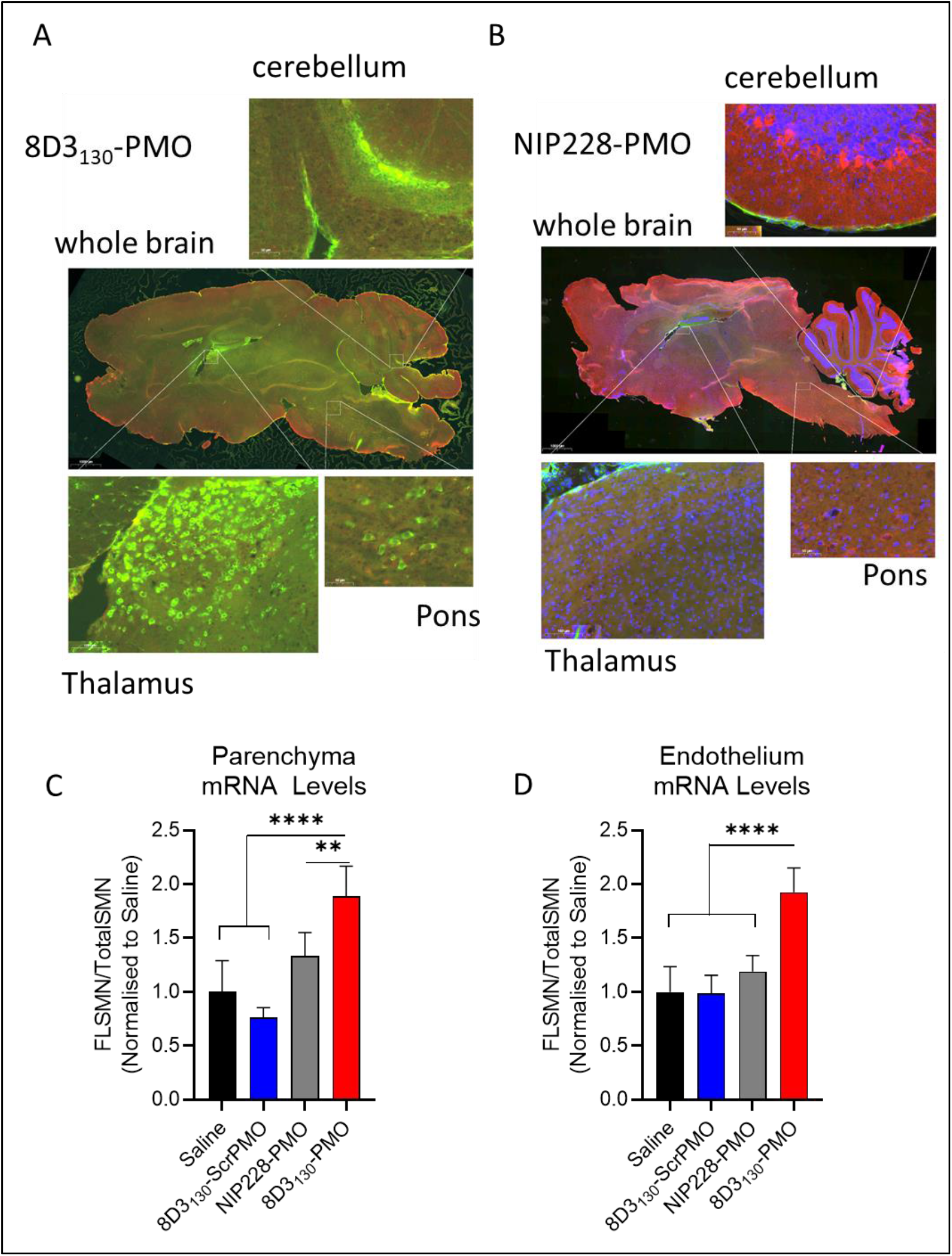
Whole brain biodistribution of antibody-ASO conjugates. Representative images of SMN2 transgenic mouse brain treated with single 50 mg/kg administration of (A) 8D3_130_-PMO or (B) NIP228-PMO. The CNS was isolated from adult mice 24 hours post administration following perfusion fixation 8D3_130_-PMO and NIP228-PMO were identified by human secondary antibody (IgG(H+L). Whole brain slides were imaged at 20x on 3D Histec pannoramic250 slide scanner. Images represent n = 3 mice. The greatest level of 8D3_130_-PMO uptake into the brain was observed in the thalamus, pons and cerebellum regions of the brain. *FLSMN2* expression via QPCR was analysed in endothelium (blood brain barrier) and parenchyma of the brain fractionated by EC extraction. Mice were treated with 50 mg/kg 8D3_130_-PMO, NIP228-PMO, 8D3_130_-scrPMO or 0.9% saline. Seven days post administration brain was collected and the endothelial cells were isolated from the brain parenchyma. (c) QPCR analysis of *FLSMN2* mRNA expression from parenchyma brain fraction. 8D3_130_-PMO was significantly more active (1.89 + 0.28) than NIP228-PMO (1.3 + 0.2), 8D3_130_-ScrPMO (0.76 + 0.09), or saline (1 + 0.3) p < 0.0001. (d) QPCR analysis of *FLSMN2* mRNA expression in endothelium brain fraction. 8D3_130_-PMO was significantly more active (1.93 + 0.23) than NIP228-PMO (1.19 + 0.15), 8D3_130_-ScrPMO (1 + 0.17), or saline (1. + 0.24) p < 0.0001. Statistical significance (representative P values) was performed in GraphPad Prism. Data shown as the mean + s.d., n = 6 per group. Results analysed with one way ANOVA corrected for multiple comparisons using Tukey’s Test.

To validate that observed fluorescence correlates with drug activity within the brain parenchyma, we performed an endothelial brain tissue fractionation experiment on adult SMA mice dosed with 50 mg/kg (n = 6 per group) to separate the endothelium of the blood-brain barrier from the brain parenchyma. Tissues were collected 7 days post administration and, for this experiment, we included the 8D3_130_-scrambled PMO (8D3_130_-ScrPMO) as a control. Clean fractionation of the endothelial cell (EC) and parenchyma fractions was validated with qRT-PCR analysis of EC markers (Pecam1 and Vcam1), neuronal makers (b-tub III and Map2) as well as a glial marker (GFAP) (Fig. S3). *FLSMN2* mRNA expression was measured in both fractions (Fig. 5, C and D). Both 0.9% saline and 8D3_130_-ScrPMO controls yielded similar results, indicating no activity from the 8D3_130_ antibody itself. 8D3_130_-PMO was active in both the brain parenchyma (1.89 ± 0.28) and endothelial cell (1.93 ± 0.23) fractions, indicating that the 8D3_130_-PMO is active in both the parenchyma of the brain itself as well as within the BBB cell compartment. NIP228-PMO activity was not significantly different from saline or 8D3_130_-ScrPMO activity.

### 8D3_130_-PMO predominantly co-localizes to astrocytes of the spinal cord

In order to evaluate which resident spinal cord cell populations were targeted by 8D3_130_-PMO, we dosed adult *SMN2* transgenic mice with 50 mg/kg 8D3_130_-PMO or NIP228-PMO and perfused animals 24 hours later. Cryosections of spinal cord were then analysed for co-localization of the 8D3_130_-PMO and NIP228-PMO to motor neuron (using ChAT immunolabelling) and astrocyte (using GFAP immunolabelling) cell populations. TfR is found on the surface of neurons in the spinal cord and so a widespread cellular distribution of 8D3_130_-PMO was expected(*11*). Instead, however, we observed a more cell-specific uptake of 8D3_130_-PMO in the astrocytes of the spinal cord while minimal co-localization was found with motor neurons (Fig. 6). The tight association of astrocytes with BBB and BSCB regulates passage of compounds into the parenchyma. It is therefore likely that astrocytes represent the first cell population that the 8D3_130_-PMO comes into contact with on exposure to the spinal cord parenchyma.

**Figure 6:**
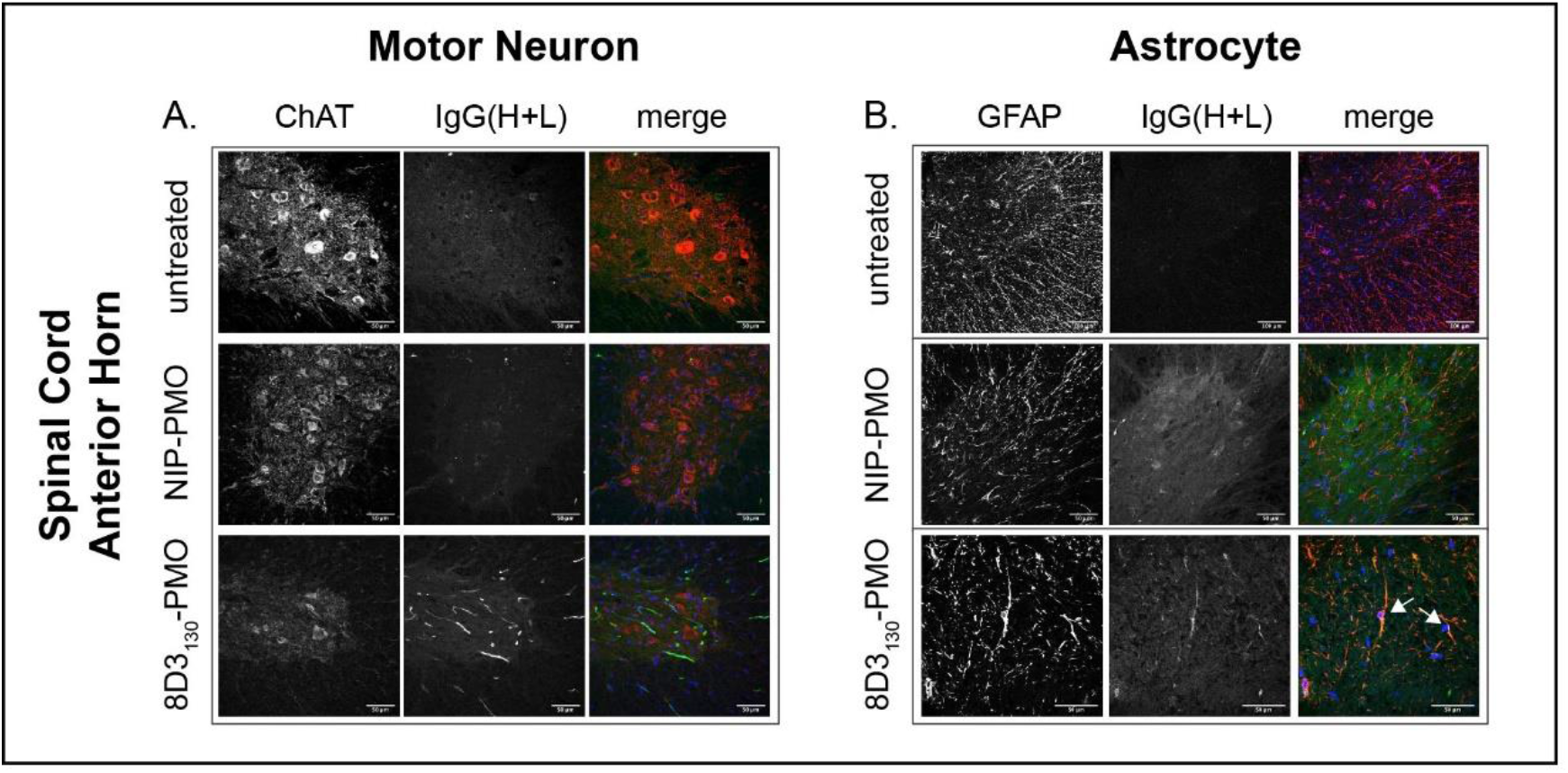
Cellular localisation of 8D3_130_-PMO to the astrocytes in the spinal cord. Representative confocal images of spinal cord following single 50 mg/kg administration of 8D3_130_-PMO and NIP228-PMO. The CNS was isolated from adult mice 24 hours post administration following perfusion fixation. (A) Motor neurons (ChAT) in the anterior horn of the spinal cord, and 8D3_130_-PMO identified by human secondary antibody (IgG(H+L)) showed no overlap (merge). Fluorescence indicated a retention of the 8D3_130_-PMO (IgG(H+L)) in the vasculature. (B) Astrocytes of the anterior grey horn (GFAP) were colocalised with 8D3_130_-PMO (IgG(H+L)) (arrow heads).

### 8D3_130_-PMO increases FLSMN2 expression and extends survival in severely affected SMA mice

The *hSMN2* mouse strain (FVB.Cg-*Smn1^tm1Hung^* Tg(SMN2)2Hung/J) can be bred to produce a severe SMA phenotype in early postnatal pups. SMA pups exhibit low body weight by day 5 and reduced movement from day 6-8 following which they typically die between 7-10 days of age (*29*). To test the potential of 8D3_130_-PMO to prevent the onset of an SMA phenotype, pups were treated with single subcutaneous administration of 20 mg/kg or 50 mg/kg 8D3_130_-PMO, NIP228-PMO or saline at post-natal day 0 (PND0). Survival and tissue expression of SMN2 mRNA was evaluated (Fig. 7). Median survival following single subcutaneous administration of 20 mg/kg 8D3_130_-PMO (n = 7), NIP228-PMO (n = 15), 8D3_130_-scrambled PMO (n = 11) or 0.9% Saline (n = 17) was 24, 12, 11 and 7 days respectively. Survival of 8D3_130_-PMO treatment was statistically greater than NIP228-PMO or 8D3_130_-scrPMO, p<0.0001 (Fig. 7A). Pups treated with the higher dose of 50mg/kg 8D3_130_-PMO (n = 7) or NIP228-PMO (n=8) survived a median of 22 and 21 days respectively (p=n.s.) (Fig. 7B). Only at the lower treatment dose was the difference in activity between 8D3_130_-PMO and NIP228-PMO evident. Activity of the NIP228-PMO compound was not unexpected, given its earlier observed activity in peripheral tissues (Fig. 4) and given that the blood-CNS tissue barriers in early post-natal day mice are developmentally less mature than in adults, even allowing the passage of non-conjugated ASO ^9^. *FLSMN2* mRNA expression was measured in a separate group of pups treated at PND0 with 50mg/kg 8D3_130_-PMO, NIP228-PMO, 8D3_130_-Scrambled PMO or 0.9% saline. Brain, spinal cord, skeletal muscles from the hindlimbs, heart, kidney and liver tissues were collected PND7. In all tissues tested, both 8D3_130_-PMO and NIP228-PMO significantly enhanced *FLSMN2* levels over saline and 8D3_130_-scrambed PMO treated tissues (Fig. 7, C to H). 8D3_130_-PMO produced greater *FLSMN2* expression over NIP228 within brain (3.5 ± 0.84 vs 2.3 ± 0.6 fold change), spinal cord (5.1 ± 1.24 vs 3.4 ± 1.2 fold change) and heart (3.8 ± 0.24 vs 2.96 ± 0.24 fold change) (Fig. 7, C, D and F).

**Figure 7:**
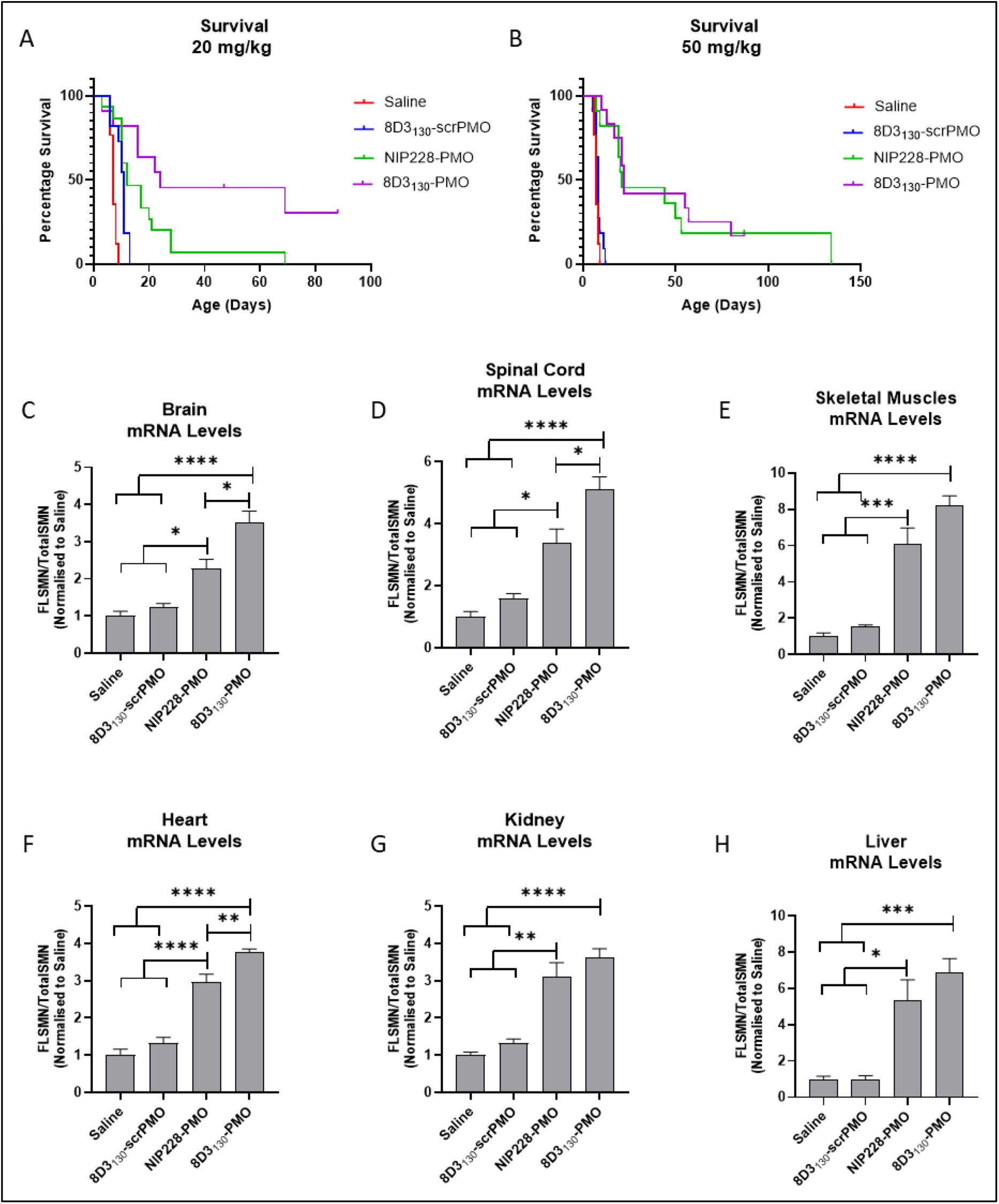
Survival and mRNA levels in severe SMA Pups treated with antibody-PMOs. (A) Survival following single subcutaneous administration of 20 mg/kg 8D3_130_-PMO (n = 7), NIP228-PMO (n = 15), 8D3_130_-scrambled PMO (n = 11) or 0.9% Saline (n = 17). Median survival was 24, 12, 11 and 7 days respectively. Mean survival after treatment with 8D3_130_-PMO was significantly greater than NIP228-PMO, p<0.0001. Stats determined by Log-Rank (Mantel-Cox) test. (B) Survival following single subcutaneous administration of 50 mg/kg 8D3_130_-PMO (n = 12), NIP228-PMO (n = 11), 8D3_130_-scrambled PMO (n = 11) or 0.9% Saline (n = 17). Median survival was 22, 21, 8 and 7 days respectively. Both 8D3_130_-PMO and NIP228-PMO was statistically significant from 0.9% saline treated group, p < 0.0001. However, there was no statistical difference between 8D3_130_-PMO and NIP228-PMO. Stats determined by Log-Rank (Mantel-Cox) test. (b-g) qRT-PCR measure of mRNA from tissues treated with 50 mg/kg antibody-PMO and collected 7 days post administration. Results were normalised to saline treatment controls. *FLSMN2* mRNA represented at ratio to total *SMN2* transcripts. One-Way ANOVA with Tukey’s multiple comparison test. All data represent mean values ± S.D. of two replicates. P-value representations: p**** < 0.001, p***<0.005, p** <0.005, p* <0.05.

## Discussion

There are currently over 80 antibody drugs approved by FDA. The majority of Ab-based drugs treat immune-mediated diseases and various cancers including hematologic malignancies and solid tumours. Only nine of the 80 approved Ab drugs are antibody-drug conjugates (ADCs), all of which are approved for cancer therapies(*37*). These ADCs act by internalizing cytotoxic small molecules into cells expressing cancer cell membrane proteins such as CD30 and CD33. Antibody-ASO conjugates are a newer class of drug only recently developed for therapeutic application(*38*).

ASO therapies have become one of the most promising forms of gene therapies for a wide range of diseases. In their naked form, ASOs are unable to pass through the BBB or BSCB and therefore require invasive modes of delivery through either direct brain injection or intrathecal administrations to treat neurodegenerative diseases. The highly successful nusinersen, an ASO targeting *SMN2* in patients with SMA, has extended survival and welfare for many children(*8*). However, the repeated intrathecal administrations required for treating neurodegenerative diseases subject patients to a lifetime of this risky procedure. Clearly, reaching the CNS via systemic administration would be a major step forward in ASO therapies. We have previously used peptides to deliver ASOs to the brain and spinal cord at therapeutically relevant levels(*24*). Here we make use of the natural mechanisms for translocation across the BBB by targeting the TfR with an anti-mouse TfR antibody 8D3_130_. Strikingly, systemic administration of 8D3_130_-PMO conjugates results in high levels of CNS exposure in an adult SMA mouse model and rescues survival in the severe SMA model.

TfR1 is a 97-kDa type 2 membrane protein expressed as a homodimer(*39, 40*). The TfR1 binds to iron-laden transferrin and translocates it across brain endothelial cells. Using an anti-TfR1 antibody to translocate a cargo across the BBB has been previously successfully carried out with various cargos(*10*). Additionally, delivery of ASOs conjugated to anti-transferrin antibody have been used to image gene expression in rat models of brain ischemia and brain glial tumors(*41–43*). However, studies into antibody-delivery of ASOs to the brain are limited. These early studies did not look into the cellular biodistribution of the anti-TfR conjugates nor the activity of the ASOs. The lack of in-depth imaging leaves doubts into the transport of the anti-TfR-conjugates across the brain endothelium and cell-specific uptake.

8D3_130_ was developed from the parent antibody 8D3, which has a stronger binding affinity to the mTfR, 130nM and 1.2nM respectively(*15*). 8D3 is capable of transducing across the BBB into the brain parenchyma and has been used as a fusion protein with erythropoietin (EPO), a glycoprotein cytokine with neuroprotective effects in mouse model of Alzheimer disease(*44*). As a fusion protein there is no linker between the 8D3 antibody and EPO. Instead, EPO is expected to bind to the extracellular EPO receptor. The 8D3-EPO fusion antibody garnered a modest effect in the AD mouse model(*45*).

8D3_130_-PMO conjugate modified the PK and activity of both the 8D3_130_ and PMO. PMO reduced the amount of 8D3_130_ assayed in the brain but only reduced 8D3_130_ in spinal cord 96 hours post-administration. Plasma levels of 8D3_130_ were also reduced when conjugated to PMO. However, 8D3_130_-PMO greatly improved uptake and activity of the PMO into the brain and spinal cord. To observe the effect of 8D3_130_ delivery of PMO to the brain and spinal cord we chose an adult mouse model with four copies of the human SMN2 transgene(*30*). This animal has no phenotype or observed disruption of the BBB or BSCB and should therefore recapitulate the biodistribution required for treating neurodegenerative and neuromuscular diseases. A two-fold change in *FLSMN2* expression is therapeutically relevant for alleviating SMA disease pathology. Here we have shown the 8D3_130_-PMO conjugates reaches 2-fold and above changes to *FLSMN2* expression in the brain and spinal cord following single dose administrations of 50 mg/kg and above. Similar levels of *FLSMN2* expression in the same mouse model have only been achievable by direct brain administrations of 100-200 ug PMO(*46*). Therapeutically relevant levels of *FLSMN2* expression were also observable in skeletal muscles, tibialis anterior and quadriceps, as well as the liver while lower levels of expression was achieved in the kidneys. In addition, doses as high as 100 mg/kg had no observable negative effects on the mice. This could be due to the natural path of kidney circulation for mAb. The majority of mAbs are reabsorbed in proximal tubules and re-enter the systemic circulation, thus mitigating the kidney toxicity seen in ASO treated mice(*47*). It remains to be seen if the 8D3_130_-PMO conjugate behaves in the same fashion.

The BBB is not a homogenous tissue and difference in permeability to the BSCB may allow for the targeted passage in the spinal cord(*48*). In addition, there are even a different levels of permeability between the sections of the spinal cord that can account for heterogenous patterning of 8D3_130_-PMO across the cervical, thoracic and lumbar regions of the spinal cord(*49*). Immunohistochemistry of the 8D3_130_-PMO compounds indicate a particularly high level of uptake into pons and thalamus regions of the brain. A similar observation was made with another anti-TfR drug conjugate, JR-141. JR-141 is an anti-human TfR-human iduronate-2-sulfatase (IDS) protein conjugate generated to treat the lysosomal storage disease mucopolysaccharidosis II (MPSII). Preclinical studies in mice and monkeys given IV administration observed JR-141 within Purkinje cells of the cerebellum and pyramidal cells in the hippocampus(*12*). The authors also observed wide spread biodistribution of JR-141 in the heart, kidney, liver, lung and spleen. However, they did not use an Ab isotype control so it is unclear if this is specific for TfR binding or due to a more general Ab uptake. These results lead to a Phase I/II clinical trial and a reduction in heparan sulfate, a lysosomal glycosaminoglycan inadequately catabolised in MPSII, was observed in the cerebral spinal fluid of treated patients(*50*).

Cell specific uptake was investigated for spinal cord delivery. A high rate of uptake was observed in the astrocyte cell population, but not in the motor neurons, of the spinal cord. Astrocytes make up a critical component of the BBB and BSCB and they are the first point of contact for compounds translocating across the endothelium. Although they themselves do not express the transferrin receptor *in vivo*, they express the Fc gamma receptor (FcγRI) that bind to the Fc component of the immunoglobulin IgG(*51*) which may facilitate translocation. Astrocytes provide metabolic support to motor neurons and low levels of SMN in astrocytes exacerbate motor neuron death in SMA(*52*). Increasing astrocyte-directed SMN expression extended survival in severe SMA line of mice(*52*) and neurons co-cultured with astrocytes cultured from SMA mice had reduced synaptic formation and transmission indicating a cell-autonomous effect in SMA derived astrocytes(*53*). Additionally, SMA motor neurons co-cultured with SMA or wild type astrocytes resulted in similar numbers of synapses and excitatory postsynaptic current, highlighting the importance of the astrocyte-motor neuron interaction in SMA(*53*). Taken together, these studies validate astrocytes as a potential target for ASO therapy in SMA.

Delivery of oligonucleotides and siRNA with antibodies has been mostly studied using the transferrin receptor as the antibody target. Early work used a system whereby an antibody is fused to avidin to act as a carrier for biotinylated pharmaceutically active drug. Penichet et al., designed a fusion anti-TfR antibody with avidin as a carrier molecule for biotinylated peptide nucleic acid (PNA) oligo(*54*). However, only 0.12% injection dose/gram was observed in brains of treated rats, far less than the 2.7% observed in our study. A similar compound, anti-TfR avidin linked to biotinylated luciferase targeting siRNA, was tested in rat brain tumour expressing luciferase and reduced luciferase levels were observed 48 hrs after IV administration(*55*). Similar to our antibody-ASO design, Sigo et al., utilized a linker to covalently conjugate the anti-TfR (antiCD71) to an siRNA(*56*). AntiCD71-siRNA compounds were only shown to be active in peripheral tissues, liver, heart and skeletal muscle. Unlike our study, delivery into the CNS was not reported and their IgG control-siRNA was not active in the skeletal muscle. The reason for this is unclear.

In addition to improved uptake into the CNS and peripheral tissues, 8D3_130_-PMO also rescued survival in a severe mouse model of SMA. A severe SMA pups carries two copies of SMN2 and deletion of exon 7 from the endogenous mSmn. These pups are born indistinguishable from littermates but begin to show reduction in weight and movement within a few days with early lethality at an average of 7 days. Due to the early onset of a disease phenotype, treatment of PMO is required within a day or two after birth. The BBB and BSCB is immature at this early stage of development, leaving the brain and spinal cord exposed high molecular weight molecule like PMOs. It was unclear if the conjugation of a PMO to 8D3_130_ would improve uptake over NIP228-PMO control. Indeed, using a high dose of 50mg/kg, both 8D3_130_-PMO and NIP228-PMO rescued survival and tissue expression of full length SMN2 mRNA. However, a significant improvement in survival of 8D3_130_-PMO over NIP228-PMO was observable using a lower dose of 20 mg/kg, indicating the TfR is active in BBB transport at these early developmental stages and the BBB is acting as a barrier, albeit a weak on(*57*). The BBB and BSCB represent a significant barrier to the delivery of biologic drugs, both protein and nucleic acid based. Improving CNS exposure using anti-TfR antibodies, which allow transcytosis across the BBB, can be exploited for the delivery of drugs to the brain. We have found that systemic dosing of an anti-TfR antibody-PMO conjugate can access the central compartment and affect the splicing of the *SMN2* mRNA to levels only previously observed with direct brain drug administration, rescuing survival in severe SMA mice. This study provides a proof of concept for therapeutic, systemic dosing of Ab-ASO conjugates to treat brain disorders and offers the hope of improved therapies for many debilitating neurological diseases.

## Materials and Methods

### Preparation of antibodies

The rat anti-mouse transferrin receptor (TfR) antibody 8D3 was mutated to reduce its affinity for mouse TfR to generate 8D3_130_ as described(*15, 23*). 8D3_130_ and a negative isotype-control antibody against the hapten nitrophenol (NIP228) were expressed as chimeric human IgG1 molecules with a cysteine residue inserted in the CH2 domain of the heavy chain to allow site specific conjugation of the PMOs(*58*). Antibodies were expressed in transiently transfected Chinese hamster ovary (CHO) cells in serum-free medium as described previously(*59*). Cultures were maintained in a humidified incubator at 37°C, 5% CO2 for 14 days after which the media was harvested. Antibodies were purified from the medium using protein affinity chromatography followed by size exclusion chromatography. The concentration of IgG was determined by A280 using an extinction coefficient based on the amino acid sequence of the IgG(*60*).

### Conjugation of thiol-derivatized antibodies with Mal-C3-FB[RB]_6_-PMO

A 25 mer maleimide-functionalized phosphorodiamidate morpholino oligo (PMO) conjugate targeting the ISS-N1 site of the SMN2 gene was synthesized by conjugation of the 3’-end of the PMO to the C-terminal carboxylic acid moiety of the linker through amide coupling (Fig. S1)(*61*). Sequence for ISS-N1 targeted PMO (5’-3’) was GTAAGATTCACTTTCATAATGCTGG and sequence for scrambled PMO (5’-3’) was CCTCTTACCTCAGTTACAATTTATA, both were fully modified phosphorodiamidate morpholino oligos.

lgG (15 ml at 10 mg/ml, ~1 μmol in PBS) was reduced with 40 eq. of TCEP (0.5M in water, 80 μl, 40 μmol) for about 3 hours at room temperature under mild agitation. Samples were taken at different time points and analysed by MALDI-TOF mass spectrometry to follow the reduction reaction. Afterwards, the buffer was exchanged to DPBS containing 1 mM EDTA by using a HiPrep 26/10 desalting column at a flow rate of 10 ml/ml to remove unreacted TCEP. The reduced antibody-containing fractions (~20 ml) were combined and re-oxidized with 400 μl of a 50 mM (20 μmol, 20 eq.) solution of dehydroascorbic acid in DMSO for 4 hours at room temperature. Subsequently, the reaction mixture was desalted three times and concentrated by ultrafiltration (Ultracel 100 kDa Ultrafiltration disc with 100 mM phosphate buffer, pH 6.9, Merck Millipore, Burlington, MA, USA) to a final volume of 12 ml.

After determination of protein concentration, a 5-fold excess of PMO-maleimide conjugate in 100 mM PBS buffer (pH 6.9) was added. After overnight incubation, the conjugation reaction was purified by size exclusion chromatography (GE HiLoad 26/600 superdex 200pg column at a flow rate of 2 ml/min PBS). Product containing fractions were combined and concentrated by ultrafiltration (Ultracel 100 kDa Ultrafiltration disc, Merck Millipore) to 10 mg/ml in DPBS.

### Antibody concentrations in mouse brain, spinal cord, and plasma via MesoScale Discovery (MSD) assay platform

Male C57B/6 mice, age 10–12 weeks were intravenously injected with anti-TfR antibody (8D3_130_) or control IgG (NIP228) with or without PMO at 20 mg/kg (2 mg/ml on DPBS) or molar equivalent. Intravenous doses were administered into a tail vein at a constant dose volume of 10 ml/kg. Antibodies were supplied in D-PBS (Sigma). Following dosing, blood plasma samples were collected from the lateral tail vein (ca 200 μL) into a Li-Hep microvette (BD Diagnostic Systems, Franklin Lakes, NJ, USA) from each of six animals per time point per dose group. A second sample (ca. 600 μL) was collected by cardiac puncture under isoflurane anaesthesia into a Li-Hep microtainer (BD Diagnostic Systems). Following collection, blood samples were allowed to clot for 30 min and centrifuged at 10,000 rcf for 2 min at 4°C and the resultant plasma collected and flash frozen on dry ice for subsequent measurement of antibody concentration.

After final blood collection the mice were perfused with DPBS at a rate of 2 ml/min for 10 min until the extremities appeared white. The spinal cord and brain were removed, weighed, and homogenized in 5 volumes of ice-cold PBS containing 1% NP-40 and Complete® protease inhibitor cocktail tablets (Roche Diagnostics, Basel, Switzerland) using 2×10 clockwise strokes with 5 s rest time. Homogenates were rotated at 4°C for 1 h before centrifugation at 13,000 g, 4°C for 20 min. The supernatant was collected for measurement of antibody concentration. In life phase and sample preparation were performed by Pharmaron (Rushden, UK).

Antibody concentrations in mouse plasma and brain- and spinal cord homogenates were measured using the MesoScale Discovery (MSD) assay platform (Meso Scale Discovery, Rockville, MD, USA). A plate-based sandwich immunoassay format where the anti-human IgG capture antibody binds to sample hIgG (± PMO), and subsequently, a specific detection antibody labelled with SULFO-TAG emits light on electrochemical stimulation. Levels of anti-TfR and control antibody in plasma, brain, and spinal cord samples were quantified by reference to standard curves generated using calibrator samples with a four-parameter nonlinear regression model. Statistical analysis was performed using an unpaired t-test in GraphPad Prism. Data shown as the mean ± standard error of the mean, n = 4 per group.

### In vivo PMO activity in SMA mouse model

The h*SMN2* transgenic mouse (SMN2, FVB.Cg-Smn1^tm1Hung^Tg(SMN2)2Hung/J) was generated as previously described(*29, 30*) and maintained at the Biomedical Sciences Unit, University of Oxford. Handlings of Tg(SMN2)2Hung/Tg(SMN2)2Hung mice (*SMN2* offspring that carry the wild-type mouse *Smn1* gene) were carried out according to procedures authorized by the UK Home Office under the Animal [Scientific Procedures] Act 1986. *In vivo* dosing studies were performed in mice at 8–9 weeks of age. 8D3_130_-PMO and NIP228-PMO were diluted in 0.9% saline and given at a volume of 10 ml/kg body weight. Seven days post administration animals were culled via rising CO_2_ and tissues were collected, flash frozen in liquid nitrogen, and stored at −80°C until analysed. Each dosage group had n = 5 mice per group with exception of 20 mg/kg dose where n = 4 mice.

### RNA extraction and qRT-PCR

RNA extraction from harvested tissues was carried out using a Maxwell® RSC simplyRNA Tissue Kit (Promega) and cDNA generated using ABI High Capacity cDNA Reverse Transcription Kits (Applied Biosystems, Waltham, MA) following manufacturer’s instructions. For skeletal muscle, a 10 min, 55°C incubation of homogenised tissue was added prior to addition of lysis buffer before the RNA extraction. qRT-PCR reaction using TaqMan® Fast Advanced Mastermix (Applied Biosystems) was performed and analysed on StepOnePlusTM real-time PCR system (Applied Biosystems). *FLSMN2* and *Total SMN2* transcripts were amplified using gene-specific primers (Table S1)(*62*). Significance was determined via 2-way ANOVA with Dunnett multiple comparisons using GraphPad Software (*p < 0.05, **p< 0.01, ***p < 0.001).

### Protein extraction and western blot

Protein was harvested from approximately 300 mg of flash-frozen tissue was homogenized in RIPA buffer (25 mM Tris-HCl, 150 mM NaCl, 1% NP-40, 0.5% sodium deoxycholate, 0.1% SDS, pH 7.5) with Complete mini proteinase inhibitors (Roche). A total of 20–30 μg of protein was separated on 10% Novex Tris-Glycine gels (Invitrogen) and transferred to polyvinylidene fluoride (PVDF) membranes. Total protein stain (30% methanol, 6.7% acetic acid, 0.0005% Fast Green FCF (Merck KGaA, Darmstadt, Germany)) stain was used as a loading control and imaged prior to blocking. Post-blocking, human SMN protein was probed for using anti-SMN, clone SMN-KH monoclonal IgG1 (Merck Millipore) and secondary antibody IRDye® 800CW goat anti-mouse IgG (LI-COR Biosciences, Lincoln, NE, USA). Membranes were imaged on Li-Cor Odyssey® FC imager and analysed with Image StudioTM software (LI-COR Biosciences).

### ELISA based measurements of oligonucleotide concentrations in tissues

ELISA detection of PMO concentration in tissues: ELISAs were conducted, as described in Burki U et al. (*31*), using a phosphorothioate probe double-labelled with digoxigenin (DIG) and biotin (BIO). The probe was used to detect concentrations of antibody conjugated PMOs in the tissues of treated mice. Probe sequence (5’-3’): [DIG]-- *C*A*G*C*A*T*T*A*T*G*A*A*A*G*T*G*A*A*T*C*T*T*A*C[BIO] was double-labelled with digoxigenin and biotin.

### Immunohistochemistry

*SMN2* mice (12–13 week old; Tg(SMN2)2Hung/Tg(SMN2)2Hung) were administered with a single intravenous dose of 50 mg/kg 8D3_130_-PMO or NIP228-PMO. Animals were perfused with 4% paraformaldehyde (PFA, in sterile PBS) 24 h after treatment. Brain and spinal cords were isolated and fixed in 4% PFA overnight. Tissues were then washed 4 times in 1× PBS and cryopreserved in 30% sucrose in 1× PBS for 3 days at 4°C. Tissues were frozen in O.C.T. compound and stored at −80°C. Brains were cut 20 μm thick along the sagittal axis while spinal cords were sectioned transverse. Slides were stored at −80°C before proceeding. Groups of n = 3 were used for each treatment group.

### Whole Brain Images

Slides were thawed at RT, rehydrated in PBS for 40 min, permeabilized in 0.1% Triton X (in PBS) for 10 min, washed twice 5 min in PBS and blocked overnight at 4°C in 3% BSA (in PBS). The next day, slides were incubated with Alexa Fluor 488 goat anti-human IgG(H+L) (Invitrogen) at 1:500 (in 3% BSA/PBS) for 2 h at room temperature (RT). Slides were imaged at Manchester University, Bioimaging Facility on a 3D Histec pannoramic250 slide scanner at 20X.

### IgG/Nissl Neurotrace co-staining

Slides were thawed at RT, rehydrated in PBS for 40 min, permeabilized in 0.1% Triton X (in PBS) for 10 min, washed twice 5 min in PBS and blocked overnight at 4°C in 3% BSA (in PBS). The next day, slides were incubated with Alexa Fluor 488 goat anti-human IgG(H+L) (Invitrogen) at 1:500 (in 3% BSA/PBS) for 2 h at room temperature (RT). Slides were then washed thrice 5 min in PBS before incubation with Nissl NeuroTrace 530/615 (Invitrogen) at 1:200 (in 3%BSA/PBS) for 20 min at RT.

### IgG/IBA1 and IgG/GFAP co-staining

Slides were thawed at RT, rehydrated in PBS for 40 min, permeabilized in 0.1% Triton X (in PBS) for 10 min, washed twice for 5 min in PBS and blocked overnight at 4°C in 3% BSA (in PBS). The next day, slides were incubated with rabbit polyclonal antibody to GFAP (Abcam, Cambridge, UK) at 1:5000, or with rabbit anti-IBA1 antibody (Wako, Osaka, Japan) at 1:1000, for 24 h in 3% BSA (in PBS) at 4°C. The next day, slides were washed thrice for 5 min in PBS before incubation with Alexa 594 goat-anti rabbit secondary antibody at 1:1,000 and Alexa Fluor 488 goat anti-human IgG(H+L) at 1:500 (both Invitrogen; in PBS) for 2 h at RT. Samples were imaged on Olympus FV1000 confocal microscope using Fluoview software. Minimal post-imaging processing was done with FiJi (ImageJ).

### IgG/ChAT co-staining

Slides were thawed at RT, rehydrated in PBS for 3 h and dried overnight at 4°C. The next day, slides were permeabilized and blocked by incubation for 4 h in 0.3% Triton X + 5% BSA (in PBS), then washed for 5 min in PBS. Slides were incubated with a rabbit antibody anti-ChAT (EPR 16590; Abcam) at 1:250 in the blocking solution for 48 h at 4°C. Slides were then incubated with Alexa 594 goat-anti rabbit secondary antibody at 1:750 and with Alexa Fluor 488 goat anti-human IgG(H+L) at 1:500 (both Invitrogen; in PBS) for 2 h at RT.

All slides were washed three times for 5 min at RT and dried before mounting with the DAPI-containing VectaMount Permanent Mounting Medium (Vector Laboratories, Burlingame, CA, USA), and sealed with nail varnish. Slides were stored at 4°C in the dark before imaging on Olympus FV1000 confocal microscope. All images across conditions were taken on the same day for a given staining. Microscopy images were processed minimally on FiJi (ImageJ).

### Isolation of endothelial cells

Cerebral endothelial cells (EC) from treated mice were extracted essentially as previously described(*63*). The brain was immediately extracted post euthanasia; cerebellum and olfactory bulbs removed; and the remaining brain tissue cut in half in ice cold DMEM. The brains were individually homogenized in fresh cold DMEM. After a brief spin at 1500 rcf and 4°C, the pelleted homogenate was resuspended in 18% dextran. The EC fraction was separated from the myelin/parenchyma layer with a 10 min centrifugation at 5000 rcf and 4°C. mRNA was extracted from EC and parenchyma using Maxwell RSC simplyRNA kit according to manufacturer’s instructions. Reverse transcription and qRT-PCR was performed as before. In order to ascertain the quality of the EC isolation, a series of qRT-PCRs were run on the cDNA with primers (Fig. S4 and Table S2) towards targets enriched in EC, neurons, or glial cells.

## Acknowledgments

We would like to thank the Kevin Talbot Lab for providing the protocol for ChAT immunohistochemistry as well as University of Oxford MICRON and Manchester University Bioimaging Facilities for their help to produce the immunofluorescent images.

## Funding

MRC-LMB/AstraZeneca/MedImmune Collaborative Blue-Sky Research Grant (MJG, FA)

UKRI Medical Research Council MR/R025312/1 (SMH, NA)

SMA Trust (SMH)

University of Oxford Medical and Life Sciences Translation Fund (SMH, JS)

Muscular Dystrophy UK 19GRO-PG36-0294 (JS)

Erasmus+ programme (LG)

Ecole de l’INSERM-Liliane Bettencourt Scholarship (LG)

University of Oxford Clarendon Fund (LG)

## Author contributions

Conceptualization: MG, MJAW, CW

Methodology: SMH, FA, MG, CW

Investigation: SMH, FA, MB, GT, IG, JS

Writing—original draft: SMH

Writing—review & editing: SMH, LG, NA, MJAW

## Competing interests

M.J.G. and M.J.A.W. are co-founders and equity holders in PepGen, a company developing peptide-conjugated oligonucleotides. C.W., M.B., G.T., and I.G., are employed by AstraZeneca. All other authors declare no competing interests.

