## Supplemental Material for "Systemic antibody-oligonucleotide delivery to the central nervous system ameliorates mouse models of spinal muscular atrophy"

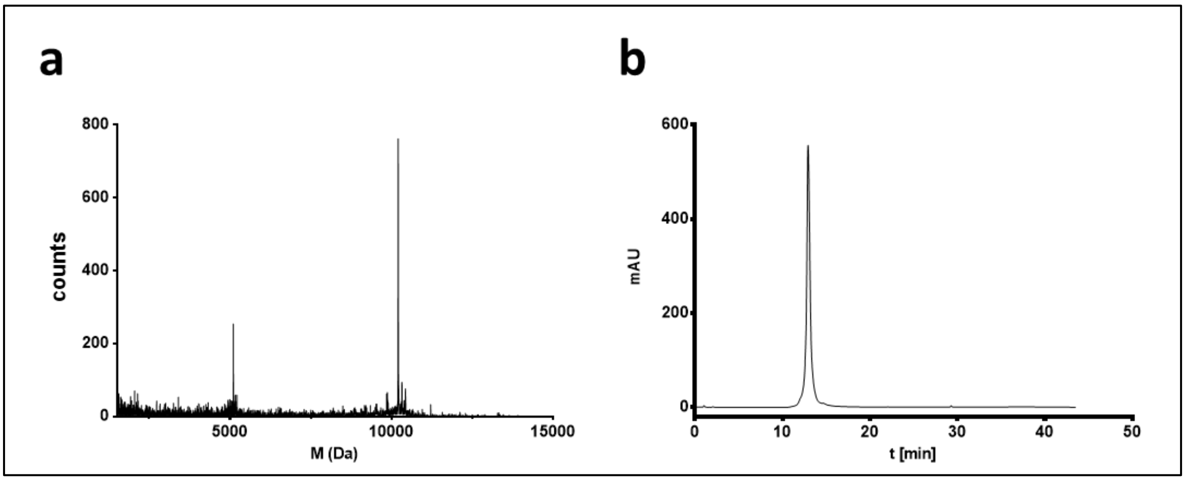


**Fig. S1.**
(A) MALDI-TOF spectra of 25-mer PMO targeting ISS-N1 and directly conjugated it to a short maleimide functionalized peptide linker, Mal-C3-FB[RB]$_6$-PMO. (B) LC-trace (260nm) of Mal-C3-FB[RB]$_6$-PMO.

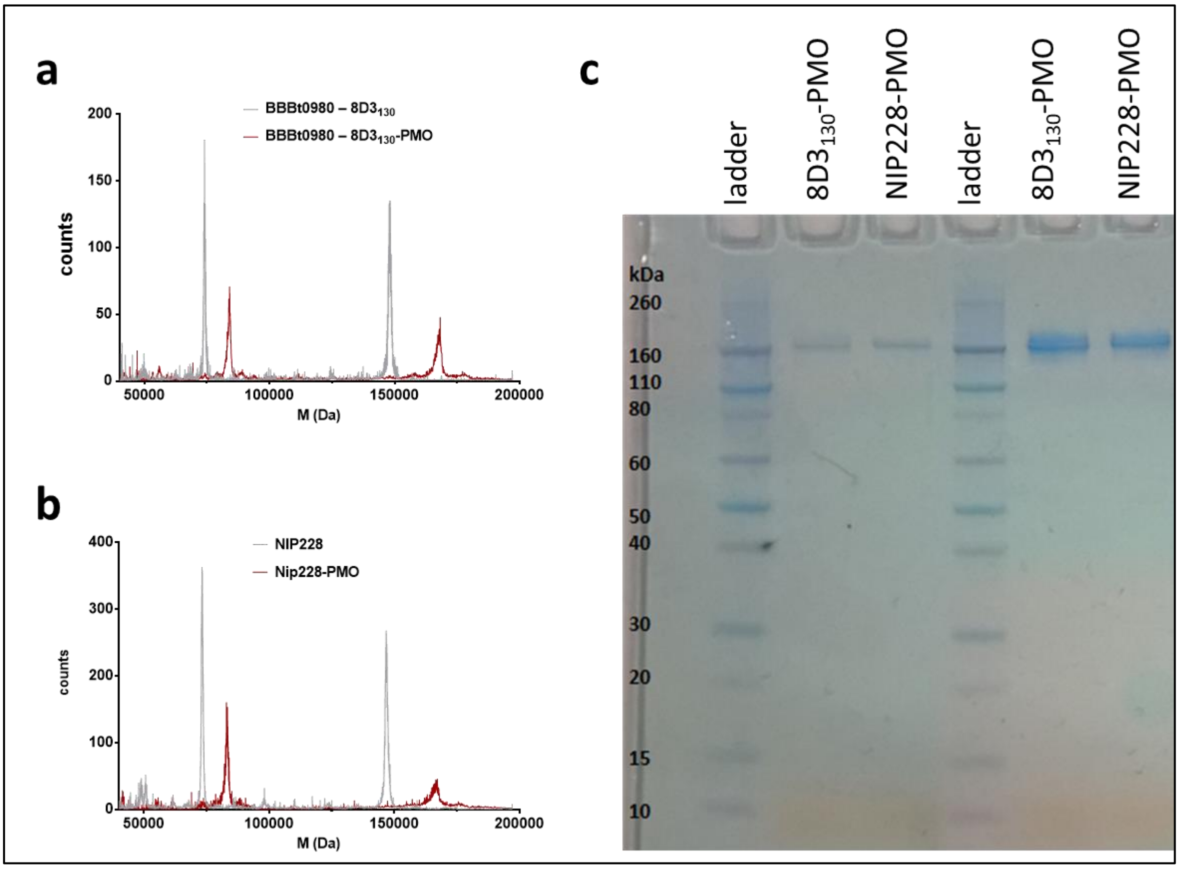


**Fig. S2.**

(A) MALDI-TOF spectra of $8D3_{130}$ before (grey line) and after (red line) conjugation reaction with Mal-PPMO. (B) MALDI-TOF spectra of NIP228 before (grey line) and after (red line) conjugation reaction with Mal-PPMO. (C) SDS-Page of antibody-PMO conjugates.

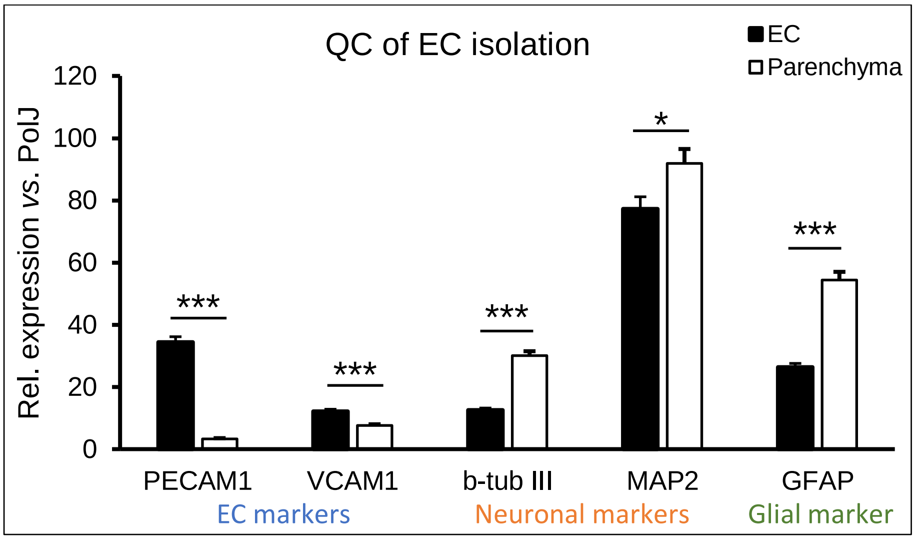


**Fig. S3.**

Endothelial cell isolation quality control data. The relative expression of endothelial cell (EC) markers, Pcam1 and Vcam1, were significantly higher in the EC fraction than the parenchyma fraction. On the other hand, the expression of neuronal markers β-tubulin III (Tubb3) and microtubule-associated protein 2 (Map2), was significantly higher in parenchyma fractions than in the EC. In addition, the expression of Glial fibrillary acidic protein (Gfap) was significantly higher in parenchyma fractions than in the EC (54.3±3.6 *vs*. 26.3±1.2) as expected. These all indicate that the EC isolation resulted in pure endothelial cells.

**Table S1.**

QPCR primers for SMN2 expression

| Primer/Probe | FLSMN2 (5’-3’) | Exon/Exon junction |
|---|---|---|
| Reverse Primer | TCGTTTCTTTAGTGGTGTCATTTAG | Ex8 |
| Forward Primer | TATCATACTGGCTATTATATGGGTTTT | Ex6-Ex7 |
| Probe | AAGGAGAAATGCTGGCATAGAGCAGC | Ex7-Ex8 |
| | **TotalSMN2 (5’-3’)** | |
| Reverse Primer | TCAGTGCTGTATCATCCCAAATG | Ex2a |
| Forward Primer | CAGGAGGATTCCGTGCTGTT | Ex1 |
| Probe | CGGCACAGGCCAGAGCGATG | Ex1-Ex2a |

**Table S2.**

QPCR primers for endothelial cell isolation quality control data

| **Primer/Probe** | ***Pecam1* (NM_008816)** |
|---|---|
| Forward | TGGTTGTCATTGGAGTGGTC |
| Probe | CACGGGTTTCTGTTTGGCCTTGG |
| Reverse | TTCTCGCTGTTGGAGTTCAG |
| | ***Vcam1* (NM_011693)** |
| Forward | GCAAAGGACACTGGAAAAGAG |
| Probe | CACTTGTGCATGGGAGACCTGTCA |
| Reverse | TCAAAGGGATACACATTAGGGAC |
| | ***Tubb3* (NM_023279)** |
| Forward | CGCCTTTGGACACCTATTCAG |
| Probe | CGCCCTCCGTATAGTGCCCTTTG |
| Reverse | TTCTCACACTCTTTCCGCAC |
| | ***Map2* (NM_008632.2)** |
| Forward | CAGGGCACCTATTCAGATACC |
| Probe | CAGCTCTCCGTTGATCCCGTTCT |
| Reverse | TCCTTCTCTTGTTCACCTTTCAG |
| | ***Gfap* (NM_010277)** |
| Forward | GAAAACCGCATCACCATTCC |
| Probe | AGACTTTCTCCAACCTCCAGATCCGA |
| Reverse | CTTAATGACCTCACCATCCCG |
